# A cortical hierarchy of sensory representations from physical structure to idiosyncratic perception

**DOI:** 10.64898/2026.07.27.741022

**Authors:** Michal M. Andelman-Gur, Tali Weiss, Lior Gorodisky, Danielle Honigstein, Ofer Perl, Edna Furman-Haran, Noam Sobel

**Affiliations:** Department of Brain Sciences, Weizmann Institute of Science, Rehovot, Israel; Psychology department, Haifa University, Haifa, Israel; Azrieli Center for Human Brain Imaging and Research, Weizmann Institute of Science, Rehovot, Israel

## Abstract

Perception begins with the physical structure of the external world, yet each of us remaps that structure into an idiosyncratic ordering shaped by individual experience and makeup. To ask how these two orderings are reflected in the brain, we used fMRI in 47 participants to measure the representation of olfactory and visual stimuli chosen such that similarities among the odorants mirrored those among the images. Olfactory and visual cortices each reflected physical similarity more strongly than idiosyncratic perceptual similarity. By contrast, a whole-brain search for the alternative uncovered the left angular gyrus, which reflected idiosyncratic perceptual similarity in both olfaction and vision. Connectivity analysis depicted this region as a likely hub for such representation, and dynamic causal modeling supported this observation. These findings reveal a transition from stimulus-centered representations in modality-specific sensory cortex to a modality-general observer-centered representation in the angular gyrus, suggesting a potentially fundamental hierarchy in brain organization.

## Introduction

Perception generates a reflection of the external world. This reflection is in part shaped by the physical features of a stimulus, but also in part remapped as a function of idiosyncratic experience and makeup^1^. A well-known example for this is the blue dress illusion, where some experience its colors as blue and black, while others see it as white and gold^2,3^. The retinal image is identical, yet stable perceptual differences emerge across observers. Where and how in the brain does such stable remapping from physical input to subjective perception occur?

To address this question we used carefully selected sets of stimuli from olfaction and vision, and described perception in a way that does not depend on words. Rather than verbal labels such as blue or gold, we described each stimulus by how similar it is to every other stimulus in its set, that is, by a relational similarity structure^4,5^. Such structures lie at the heart of perception, and generate stimulus coordinate spaces that support recognition, generalization and associative learning^4^. For each stimulus set we then obtained two such structures: one computed from the physical structure of the stimuli themselves, and one from the similarity each participant reported experiencing. Finally, we applied representational similarity analysis (RSA), which asks whether the arrangement of neural activity patterns preserves the arrangement of the stimuli they represent^6–9^. This allowed us to ask, within the same individuals and in two senses, which brain regions order the world according to physical structure, and which order it as the observer does.

We found that olfactory and visual cortices each ordered the stimuli more as the physical structure of the stimuli themselves are ordered than as participants reported perceiving them. Sensory cortex therefore retained physical information that conscious report did not reflect. By contrast, a whole-brain search for the opposite signature converged on a single region, the left angular gyrus, that ordered the stimuli as each individual perceived them, and did so in both olfaction and vision. Connectivity analysis further depicted the left angular gyrus as a likely hub for this idiosyncratic representation, and dynamic causal modeling supported this observation. These findings reveal a transition from stimulus-centered representations in modality-specific sensory cortex to a modality-general observer-centered representation in the angular gyrus, suggesting a potentially fundamental hierarchy in brain organization.

## Results

### We generated matching stimulus-similarity structures across olfaction and vision

In Experiment 1, we selected a stimulus set of odorants and visual objects with matched within-modality physical similarity structure (see Methods). This matching is important because a combined analysis across the two senses is interpretable only if their relational geometries are equated, rather than being a mix of two unrelated structures. To obtain a single physical value for each stimulus pair, we did not measure physical olfactory space as typically defined by chemistry (e.g., carbon chain length), or physical visual space as typically defined by optics (e.g., luminance). Instead, in both senses we used models that link physical structure to perception within a predictive framework: angle-distance in olfaction^10,11^, and SPoSE in vision^12–14^. Although both models were originally developed using human behavioral data, once developed they return distances computed from the stimulus alone, with no observer in the loop (see Methods). These measures have predicted perceptual similarity from physical structure alone in both olfaction^10,11^ and vision^14,15^.

After fine-tuning from a larger pilot set in 30 participants, we retained five olfactory and five visual stimuli with matched physical similarity matrices (Fig. 1a,b). Next, a separate experimental group of 47 participants (age 29.3 ± 7.08 years; 25 male, 22 female) rated the perceived similarity of all 10 unique stimulus pairs in each modality three times, yielding 2,820 ratings, of which 2,679 were retained after missed trials and exclusion of inconsistent repeats (Methods). Despite the limited number of pairwise comparisons (n = 10), physical and perceived similarities were positively correlated in olfaction (Pearson r = 0.66, t(8) = 2.46, P = 0.02, Fig. 2a,d), with a similar but non-significant trend in vision (r = 0.46, t(8) = 1.45, P = 0.09; Fig. 2b,e). More importantly, and in alignment with our intentions, olfactory and visual perceived similarities were significantly correlated with one another (r = 0.63, t(8) = 2.31, P = 0.025; Fig. 2c). In other words, as intended, we generated two sets of stimuli with matched internal organization: the perceived similarity matrix amongst the odorants mirrored the perceived similarity matrix amongst the images.

**Fig. 1.**
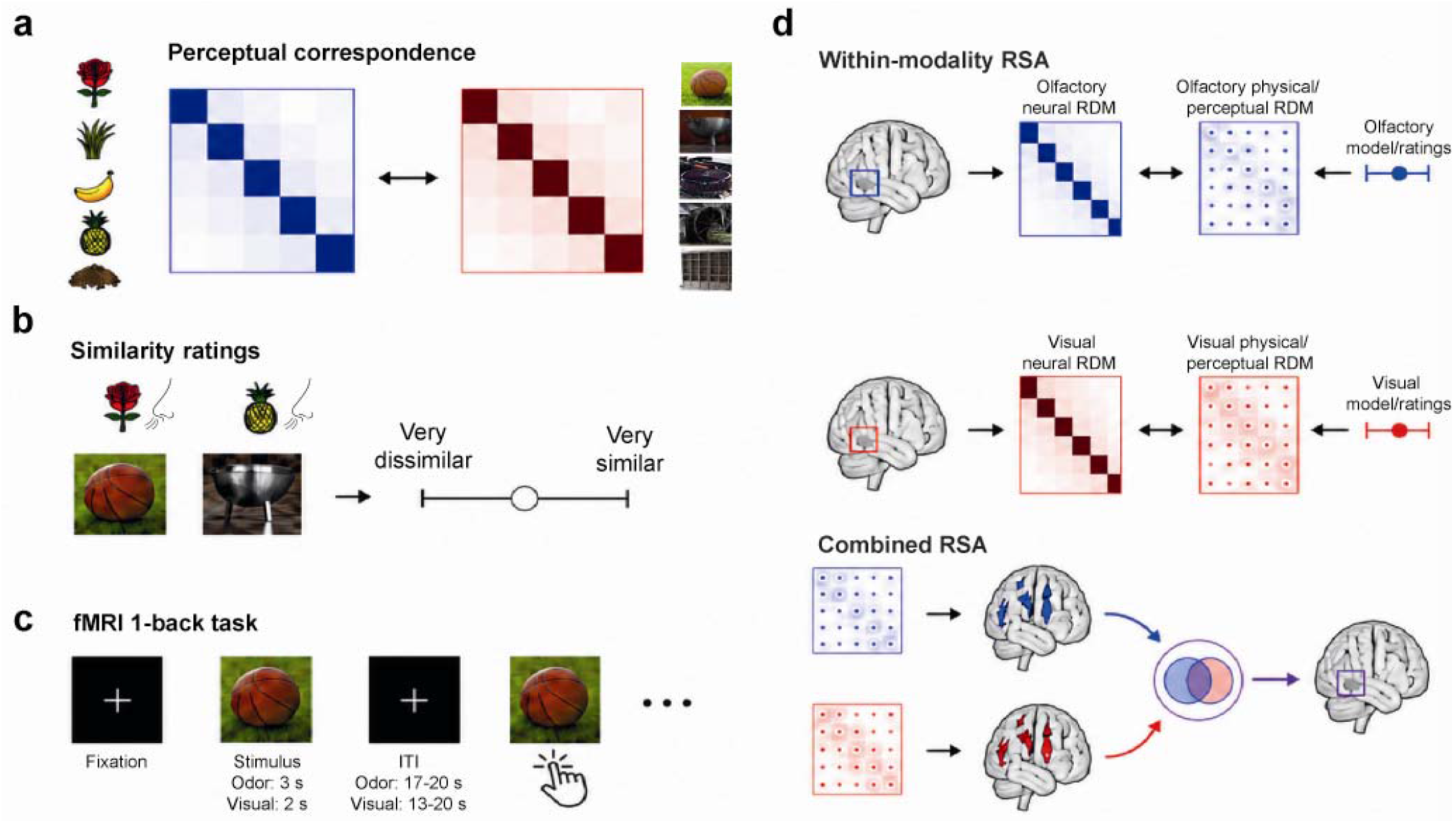
Experimental design and representational similarity analysis framework. **a,** Stimulus-selection logic. Five odorants and five visual objects were selected to maximize correspondence between their similarity structures (RDMs). **b,** In-lab behavioral similarity-rating task. Participants rated the perceived similarity of stimulus pairs within each modality. **c,** fMRI 1-back task. During scanning, participants were presented with odor or visual stimuli and responded whenever the current stimulus matched the immediately preceding stimulus. **d,** Representational similarity analysis (RSA) framework. For within-modality RSA, local multivoxel fMRI patterns were used to construct neural RDMs, which were compared with modality-matched physical and perceptual RDMs. For combined RSA, perceptual RSA maps from olfaction and vision were combined to identify shared neural representations.

**Fig. 2.**
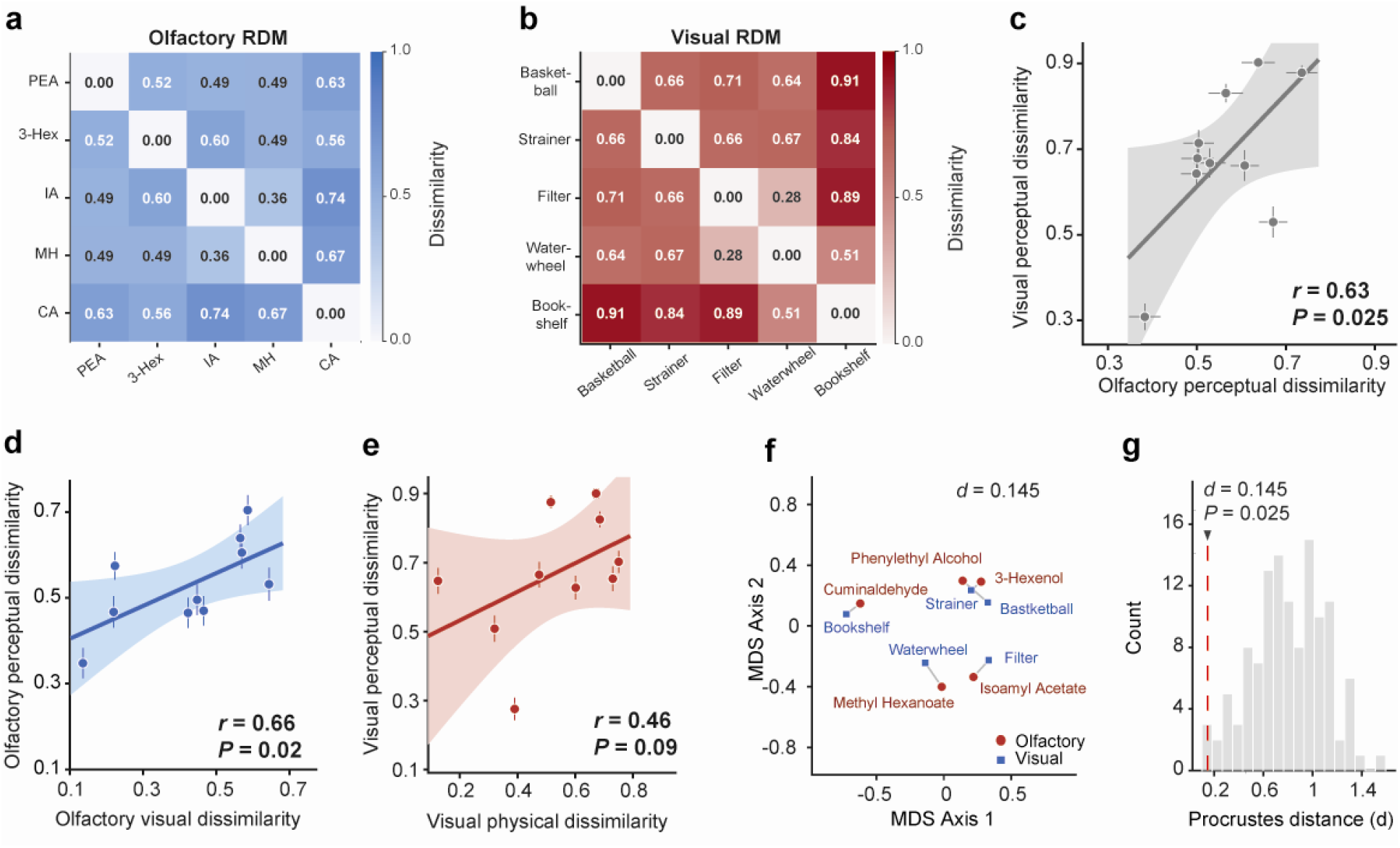
A matching stimulus-similarity structure across olfaction and vision. **a-b,** Group-mean olfactory (a) and visual (b) behavioral dissimilarity matrix (RDM) for the five odorants and five images, respectively. Similarity ratings were converted to proportional dissimilarities within participant before averaging. PEA = Phenethyl Alcohol, 3-Hex = 3-Hexenol, IA= Isoamyl Acetate, MH = Methyl Hexanoate, CA = Cuminaldehyde. **c,** Intermodal behavioral alignment between olfactory and visual perceptual dissimilarities. Each point denotes one pairwise stimulus relation from the vectorized olfactory and visual RDMs. Pearson *r* correlation and the corresponding *P*-values are indicated. Error bars indicate SEM. **d,** Correspondence between olfactory physical dissimilarity (physicochemical odor model) and group-mean olfactory perceptual dissimilarity. **e,** Correspondence between visual physical dissimilarity (SPoSE visual model) and group-mean visual perceptual dissimilarity. **f,** Multidimensional scaling (MDS) embedding of subject-level olfactory and visual perceptual dissimilarity structures. Blue circles denote odors and red squares denote visual images. Lines connect matched odor–image pairs. **g,** Procrustes permutation analysis; the histogram displays Procrustes distances obtained across all alternative odor-image assignments in MDS space; the red line indicates the observed perceptual alignment.

To visualize this relationship, we embedded the subject-level olfactory and visual dissimilarity structures into a common low-dimensional space using multidimensional scaling (MDS; Fig. 2f). In this space, the distance between two points reflects perceptual similarity: similar odors or images are placed close together, whereas dissimilar odors or images are placed farther apart. To formally quantify the correspondence, we aligned the odor and visual configurations using Procrustes analysis in the two-dimensional MDS space and compared the observed alignment with all possible odor-image assignments. Procrustes analysis asks how well one similarity map can be rotated, shifted, flipped and scaled to match another similarity map, without changing the distances among stimuli within each map. The perceptual alignment yielded a significantly smaller Procrustes distance than under alternative mappings (d = 0.145, P = 0.025; Fig. 2g), indicating that the perceptual alignment minimized the geometric mismatch between the two perceptual spaces. Notably, although the two modalities shared this group-level relational structure, individuals differed in how closely they tracked it. Similarity judgments agreed with the group consensus significantly above chance in both modalities (leave-one-out agreement: odor ρ = 0.396 [95% CI, 0.274 to 0.506], t(46) = 6.09, P = 1.1×10□□; vision ρ = 0.741 [0.679 to 0.793], t(46) = 15.17, P = 7.9×10□²□), but agreement was higher for images than for odors (paired t(46) = 6.5, P = 2.5×10□□, dz = 0.95), indicating that for this particular set of stimuli olfactory similarity judgments were more idiosyncratic across individuals (Extended Data Fig. 1).

### Olfactory and visual cortex reflected physical similarity over perceptual similarity

In Experiment 2, we next probed the neural representation of these stimuli in the same 47 participants from Experiment 1. Participants were scanned a median of 8 days after the psychophysical similarity-rating experiment (IQR = 4 – 14 days). During scanning, stimuli were presented in a 1-back task to ensure attention, and no explicit similarity judgments were made. In two olfactory runs, each odorant was delivered six times per run, yielding 60 odor trials. In the visual run, each image was presented 12 times, yielding 60 visual trials. Participants pressed a button whenever the current stimulus was identical to the immediately preceding stimulus. Performance at the 1-back task was significantly above chance in both modalities, with d′ significantly greater than zero for odors (1.5 ± 0.69 (mean ± SD), *t*_(46)_ = 14.9, *P* < 10□¹□) and images (2.5 ± 0.76 (mean ± SD), *t*_(46)_ = 22.2, *P* < 10□²□), confirming engagement in both modalities. To equate temporal alignment across modalities, and to account for any respiratory modulation of brain activity^16–21^, both odor and image presentations were triggered and time-locked to nasal inhalation^22^.

We first asked how physical and perceptual olfactory and visual similarities were reflected in olfactory and visual cortex. For olfaction, we defined four a priori regions of interest previously highlighted in the literature: anterior piriform cortex, posterior piriform cortex, orbitofrontal cortex and amygdala^23,24^. For vision, we also defined four a priori regions of interest within the visual system: V1/V2, fusiform face complex, lateral occipital cortex, and inferior temporal cortex^25–27^. For each participant, we correlated the neural RDM in each ROI with two predictor RDMs: (i) the physical (angle-distance for odorants, SPoSE for images), and (ii) the perceptual (participant’s own reported perceptual similarity). All RSA values were Fisher-z-transformed before group-level inference.

A 2×2 repeated-measures ANOVA on RSA values averaged across the four ROIs of each modality, with factors Region-Type, olfactory or visual, and Similarity-Measure, physical or perceptual, revealed a strong main effect of Similarity-Measure, *F*_(1,46)_ = 15.5, *P* = 2.8×10□□, η*²p* = 0.25. Thus, physical similarity predicted neural geometry in these areas more strongly than reported perceptual similarity. There was no main effect of Region-Type, *F*_(1,46)_ = 0.1, *P* = 0.75, and no Region-Type×Similarity-Measure interaction, *F*_(1,46)_ = 1.1, *P* = 0.3, η*²p* = 0.02, indicating a comparable pattern across olfactory and visual systems. The physical similarity advantage held within each modality-specific ROI set, both in olfactory regions (ρ = 0.214 vs. 0.104; *t*_(46)_ = 2.1, *P* = 0.02, *dz* = 0.31) and in visual regions (ρ = 0.230 vs. 0.053; *t*_(46)_ = 3.88, *P* = 1.7×10□□, *dz* = 0.57). Consistent with this effect, physical similarity RSA was significant when averaged across both olfactory ROIs (mean *ρ* = 0.214 [95% CI, 0.123 to 0.301], *t*_(46)_ = 3.9, *P* = 1.6×10□□) and visual ROIs (mean *ρ* = 0.230 [95% CI, 0.164 to 0.295], *t*_(46)_ = 5.66, *P* = 4.7×10□□), whereas reported perceptual similarity was weaker, reaching significance across olfactory ROIs (mean *ρ* = 0.104 [95% CI, 0.026 to 0.181], *t*_(46)_ = 2.24, *P* = 0.015, *dz* = 0.33) but not visual ROIs (mean *ρ* = 0.053 [95% CI, -0.024 to 0.127], *t*_(46)_ = 1.16, *P* = 0.13, *dz* = 0.17).

Planned FDR-corrected ROI-level analyses revealed that the physical odor distances significantly predicted neural geometry in all four olfactory ROIs: OFC (*ρ* = 0.31 [95% CI, 0.2 to 0.42], *t*_(46)_ = 4.45, *dz* = 0.65, *q*FDR = 2.2×10□□), anterior piriform cortex (*ρ* = 0.2 [95% CI, 0.07 to 0.32, *t*_(46)_ = 2.65, *dz* = 0.39, *q*FDR = 0.02), amygdala (*ρ* = 0.19 [95% CI, 0.03 to 0.33], *t*_(46)_ = 1.99, *dz* = 0.29, *q*FDR = 0.042), and posterior piriform cortex (*ρ* = 0.15 [95% CI, 0.01 to 0.28], *t*_(46)_ = 1.83, *dz* = 0.27, *q*FDR = 0.047; Fig. 3a). Likewise, the physical image distances significantly predicted neural geometry in all four visual ROIs: inferior temporal cortex (*ρ* = 0.29 [95% CI, 0.21 to 0.36], *t*_(46)_ = 5.95, *dz* = 0.87, *q*FDR = 1.4×10□□), LOC (*ρ* = 0.23 [95% CI, 0.13 to 0.31], *t*_(46)_ = 4.04, *dz* = 0.59, *q*FDR = 4.0×10□□), V1/V2 (*ρ* = 0.22 [95% CI, 0.1 to 0.34], *t*_(46)_ = 2.94, *dz* = 0.43, *q*FDR = 0.007), and FFC (*ρ* = 0.18 [95% CI, 0.08 to 0.29], *t*_(46)_ = 2.75, *dz* = 0.4, *q*FDR = 0.008; Fig. 3b). These results are consistent with previous findings in olfaction^23,24,28^ and vision^25–27^, indicating that physical similarity is reliably captured in modality-specific neural geometry.

**Fig. 3.**
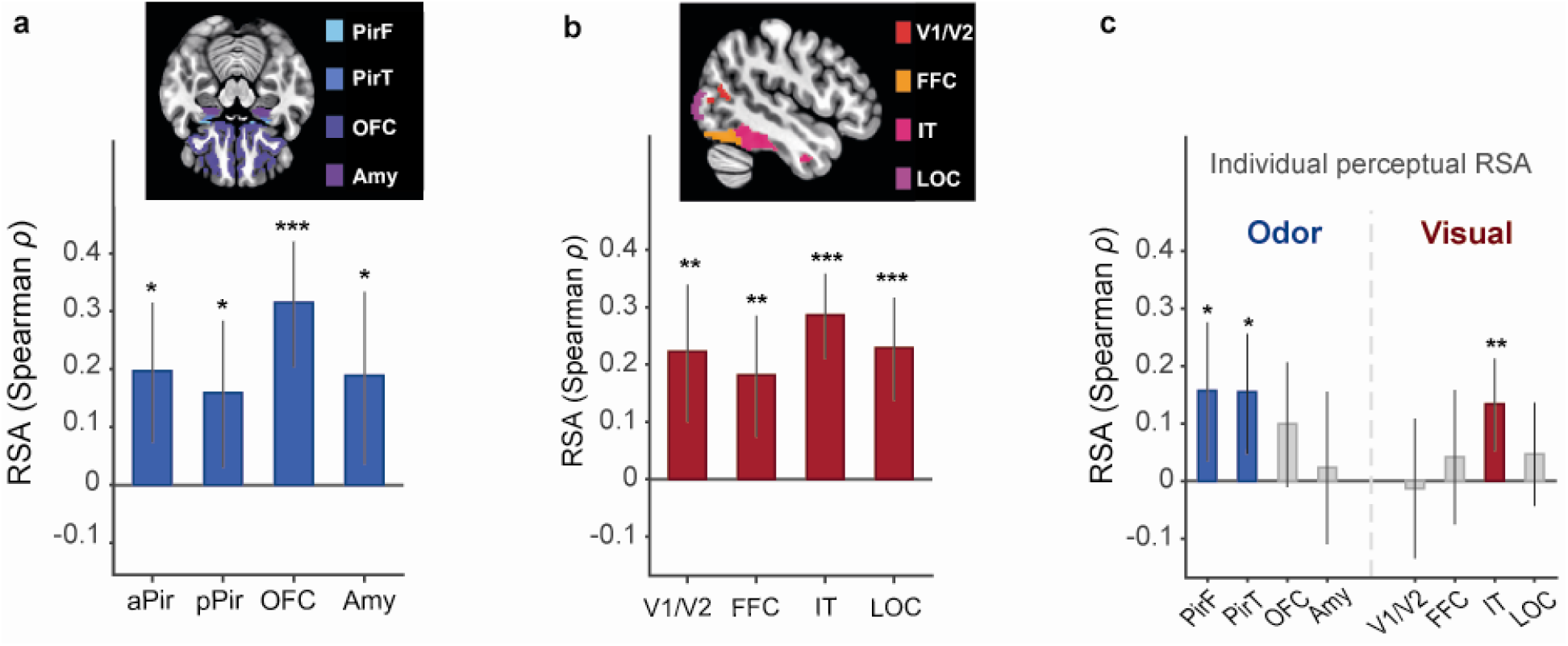
Sensory-specific neural substrates reflected sensory-specific physical similarity. **a,b,** ROI-level RSA for olfactory and visual physical models. **a,** Bars represent mean RSA between olfactory neural RDMs and the physical odor similarity RDM across olfactory ROIs. **b,** Bars represent mean RSA between visual neural RDMs and the physical visual similarity RDM across visual ROIs. Top insets present the corresponding ROI masks. **c,** Individual perceptual RSA across olfactory and visual ROIs. Bars represent mean RSA between each participant’s neural RDMs and their individual perceptual similarity RDMs. Error bars indicate SEM. Stars denote FDR-corrected significance across eight planned ROI tests (4 ROIs × 2 predictors) within each modality-specific analysis: \**q*FDR < 0.05, \*\**q*FDR < 0.01, \*\*\**q*FDR < 0.001. N = 47 participants. PirF, piriform frontal; PirT, piriform temporal; OFC, orbitofrontal cortex; Amy, amygdala; V1/V2, visual areas 1/2; FFC, fusiform face complex; IT, inferior temporal cortex; LOC, lateral occipital cortex.

Reported perceptual similarity had a more selective pattern. Planned ROI-level analyses revealed significant effects in anterior piriform cortex (*ρ* = 0.16 [95% CI, 0.03 to 0.27], t(46) = 2.15, dz = 0.31, qFDR = 0.037) and posterior piriform cortex (*ρ* = 0.15 [95% CI, 0.05 to 0.25], t(46) = 2.41, dz = 0.35, qFDR = 0.027) for odors, and in inferior temporal cortex for images (*ρ* = 0.13 [95% CI, 0.05 to 0.22], t(46) = 2.70, dz = 0.39, qFDR = 0.008; Fig. 3c). Additional control analyses are presented in Extended Data Fig. 2-3. Group-average behavioral RDMs had significant effects in OFC for odors and inferior temporal cortex for visual objects. Crossed physical-to-neural RSA revealed limited evidence for cross-modal representational structure. Odor-physical effects were observed in visual cortex, consistent with previous reports of fusiform and visual-cortical recruitment during odor identification and familiarity processing ^29–31^. By contrast, no FDR-corrected visual-physical effects were found in olfactory ROIs. This analysis suggests that the strongest similarity signals remained mostly within the appropriate sensory system, rather than appearing elsewhere in the brain. Together, these findings indicate that physical similarity provided the strongest and most consistent account of modality-specific neural geometry. Reported perceptual similarity was reflected more selectively, in piriform cortex for odors and in inferior temporal cortex for images, and even in these regions it predicted activity patterns significantly less well than the physical distances. This was not a consequence of the higher reliability of the physical measures, which control analyses ruled out (Extended Data Table 1).

### The left angular gyrus reflected perceptual similarity in both olfaction and vision

To identify regions representing reported perceptual similarity, we conducted exploratory whole-brain searchlight analyses separately for olfaction and vision using an 8-mm radius. For each participant and modality, the local neural RDM was correlated with that participant’s perceptual similarity RDM. Adaptive Storey FDR maps revealed distributed modality-specific patterns (*q*FDR < 0.05, Extended Data Fig. 4), with their largest overlap in the left angular gyrus. To ask whether this overlap survives strict correction, we searched for regions carrying perceptual similarity information across the two modalities by Fisher-z transforming each participant’s modality-specific RSA maps and averaging them across olfaction and vision^8^. The combined analysis identified corrected clusters in the left angular gyrus (221 voxels; peak combined ρ = 0.215; MNI peak: -39, -55, +29; cluster-level *P*_FWE_ = 0.015; Fig. 4a) and the right cerebellum (148 voxels; peak combined ρ = 0.193; MNI peak: +13, -79, -25; cluster-level *P*_FWE_ = 0.037). Only the angular gyrus remained significant with a 6-mm searchlight radius (*P*_FWE_ = 0.034; cerebellum *P*_FWE_ = 0.113), therefore, subsequent analyses focused on this cluster, which overlapped with the independently defined Glasser PGi parcel.

**Fig. 4.**
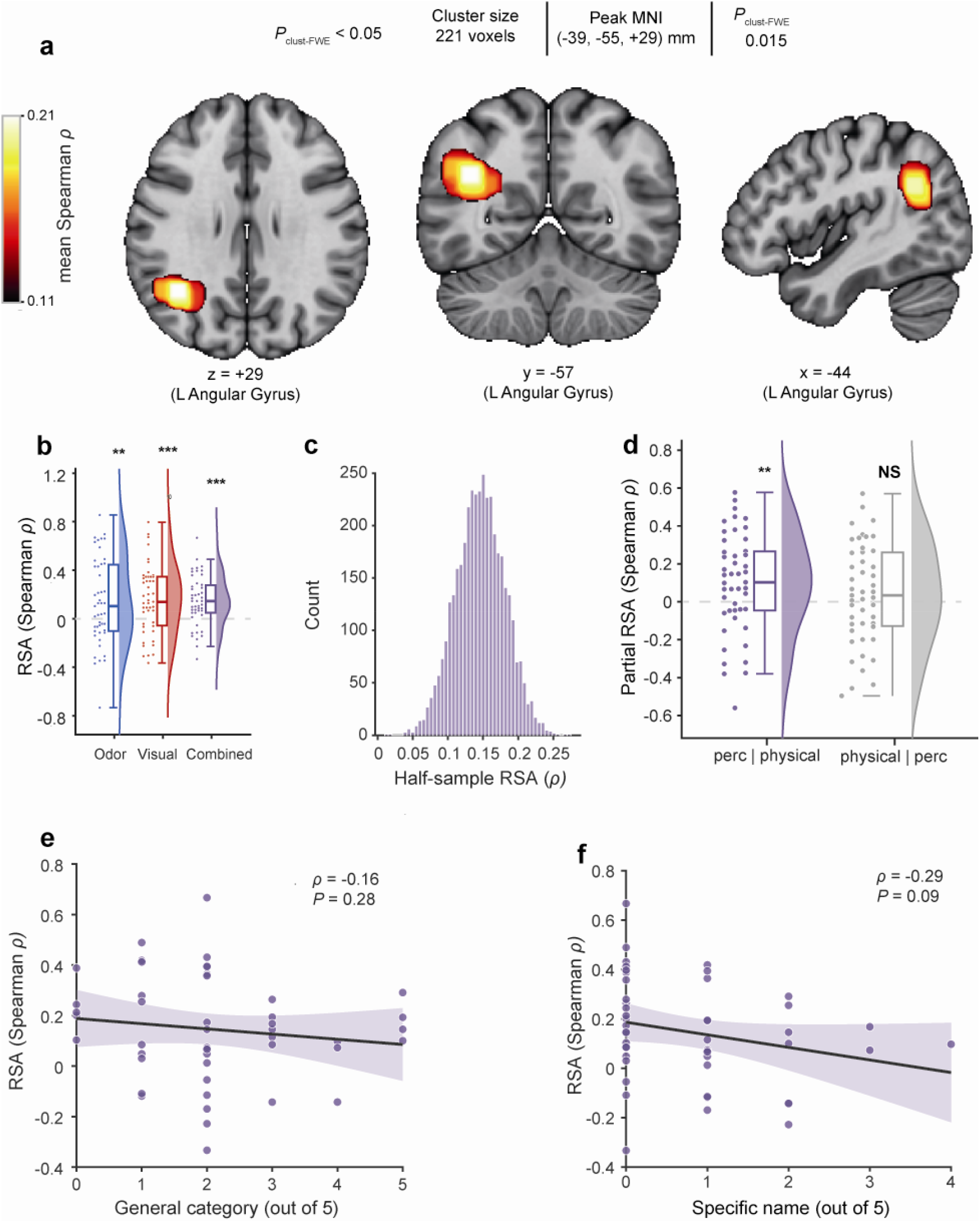
Left angular gyrus reflected perceived similarity structure in both olfaction and vision. **a,** Whole-brain searchlight RSA revealed a significant left angular gyrus cluster (8 mm radius; cluster-level FWE corrected). **b,** Mean combined, odor and visual RSA extracted from the significant angular gyrus cluster across participants. Dots represent individual participants; box and whiskers indicate median and interquartile range; violins represents distribution. **c,** Split-half robustness analysis. Distribution of mean combined RSA values obtained from repeated random half-sampling of participants. d, Intermodal perceptual alignment specificity analysis. Explained variance after controlling for physical model (physicochemical for odors, SPoSE embedding for images) or individual perception. Dots represent individual participants; box and whiskers indicate median and interquartile range; half-violin represents distribution. e-f, Scatter plots presenting Intermodal combined RSA as a function of odor category identification (e) and specific odor identification (f). 95% CI are color marked. Correlation values and *P*-vals are indicated. N = 47 participants. \**P* < 0.05, \*\**P* < 0.01, \*\*\**P* < 0.001.

To characterize the signal within this cluster, we extracted mean RSA scores for each participant. Combined perceptual RSA was significantly above zero across the group (cluster-mean ρ = 0.167 [95% CI, 0.11 to 0.22]; *t*_(46)_ = 4.74, *q*FDR = 3.1×10□□, *dz* = 0.69; Fig. 4b), with positive values in 38 of 47 participants. To determine whether the combined effect was driven by a single modality, we tested perceptual RSA separately within olfaction and vision. Perceptual RSA was significant in both modalities (odor: mean ρ = 0.162 [95% CI, 0.06 to 0.26]; *t*_(46)_ = 2.67, *q*FDR = 0.005; visual: mean *ρ* = 0.171 [95% CI, 0.09 to 0.25]; *t*_(46)_ = 3.71, *q*FDR = 4.2×10□□). Because these estimates were not independent of cluster selection, we validated them using leave-one-subject-out voxel selection within a fixed angular-gyrus search region. Both effects remained significant in held-out participants (odor: mean *ρ* = 0.135 [95% CI: 0.06 to 0.21], *t*_(46)_ = 3.15, *P* = 0.0014, *dz* = 0.46; vision: mean *ρ* = 0.101 [95% CI: 0.04 to 0.16], *t*_(46)_ = 2.87, *P* = 0.003, *dz* = 0.42), each exceeding a spatially matched random-voxel null (*P* < 0.001; 0/1,000 permutations exceeded the observed value). The modality-specific effects were robust to the number of selected voxels and leave-one-pair-out analyses, and the combined effect remained positive across all random half-splits of participants (all modality-specific *P* ≤ 0.02; Fig. 4c). These cross-validated results confirmed that both olfactory and visual perceptual similarity contributed to the angular-gyrus effect.

We next asked whether the angular gyrus signal reflected perceived similarity beyond physical similarity. Visual physical similarity was represented within the cluster (*ρ* = 0.207, *q*FDR = 5.4 × 10□□), whereas olfactory physical similarity was not (*ρ* = 0.039, *q*FDR = 0.29). We therefore tested whether perceptual similarity explained neural structure beyond the corresponding physical model using partial RSA. Perceived similarity explained unique variance beyond physical similarity (perceived given physical: mean partial *ρ* = 0.105 [95% CI: 0.029 to 0.179]; *t*_(46)_ = 2.79, *q*FDR = 0.008; physical given perceived: mean partial *ρ* = 0.057 [95% CI: -0.022 to 0.135]; *t*_(46)_ = 1.44, *q*FDR = 0.08; Fig. 4d). Because measurement noise attenuates perceptual RSA, the lower reliability of the perceptual RDM cannot explain its advantage over the noise-free physical model and, if anything, biases the comparison against perception. Thus, the angular gyrus represented perceived similarity in both modalities and retained perceptual information beyond that captured by physical similarity.

To ask whether this result reflects a genuine dissociation from the previous ROI analysis, we compared the physical-minus-perceptual RSA difference between the a priori sensory ROIs and the angular gyrus. This difference was significantly reduced in the angular gyrus in olfaction (sensory, 0.112; angular gyrus, -0.124; difference = 0.237 [95% CI: 0.08 to 0.40]; t(46) = 2.98, P = 0.005), vision (sensory, 0.181; angular gyrus, 0.037; difference = 0.145 [95% CI: 0.00 to 0.29]; t(46) = 2.02, P = 0.049) and across modalities (difference = 0.191 [95% CI: 0.07 to 0.31]; t(46) = 3.13, P = 0.003). This dissociation was confirmed using leave-one-subject-out cross-validated angular-gyrus estimates in olfaction and across modalities (olfactory difference = 0.209 [95% CI: 0.06 to 0.36], t(46) = 2.77, P = 0.008; combined difference = 0.141 [95% CI: 0.03 to 0.26], t(46) = 2.47, P = 0.017). Thus, sensory cortex was more strongly aligned with physical stimulus structure, whereas the angular gyrus was more strongly aligned with reported perceptual structure.

We next asked whether the angular gyrus better reflected individual or shared perceptual structure. We repeated the analysis using a leave-one-participant-out consensus RDM that never included the tested participant’s own ratings. For images, the leave-one-out consensus RDM was itself significantly encoded, and individual ratings did not outperform it (mean *ρ* = 0.157 [95% CI, 0.039 to 0.270], *t*_(46)_ = 2.67, *dz* = 0.39, *P* = 0.0052; paired individual vs consensus: *t*_(46)_ = 0.30, *dz* = 0.04, *P* = 0.77). For odors, by contrast, the consensus RDM was not significantly encoded (mean *ρ* = -0.060 [95% CI, −0.174 to 0.056], *t*_(46)_ = -1.04, *P* = 0.30), and individual ratings significantly outperformed it (paired individual vs consensus: *t*_(46)_ = 3.49, *dz* = 0.51, *P* = 1.1×10□^3^). Thus, especially for odors, the angular gyrus tracked each individual’s perception rather than the group-average similarity structure.

We next tested alternative explanations for the angular gyrus effect. To ensure the angular-gyrus perceptual effect did not merely reflect low-level odor attributes, we built dissimilarity matrices from pairwise differences in rated pleasantness and intensity (*N* = 44 with complete ratings; Extended Data Fig. 6). Neither attribute was encoded in the cluster (pleasantness *ρ* = 0.02, *P* = 0.75; intensity *ρ* = 0.05, *P* = 0.54), and perceptual RSA remained unchanged after controlling for both (unadjusted *ρ* = 0.21; adjusted *ρ* = 0.20; *t*_(43)_ = 3.13, *P* = 0.003). To test semantic mediation, we constructed a lexical-semantic RDM for each modality from WordNet taxonomic similarity^32^ of the image/odor source labels and correlated it with the angular-gyrus neural RDM. Semantic structure was not significantly encoded (combined mean *ρ* = 0.07, *t*_(46)_ = 1.60, *P* = 0.12; odor *ρ* = 0.08, *P* = 0.20; visual *ρ* = 0.07, *P* = 0.24) and was uncorrelated with perceived similarity (*ρ* = 0.04 and -0.02). Consistent with this, combined RSA was not positively associated with odor identification: category accuracy (*ρ* = -0.16, *qFDR* = 0.281) and specific naming were unrelated to the combined individual RSA (*ρ* = −0.29, *qFDR* = 0.096; Fig. 4e,f). Visual identification was near ceiling, precluding an equivalent analysis. Together, these results argue against explanations based on basic odor attributes or shared verbal-semantic structure.

### The left angular gyrus is the hub of a network reflecting idiosyncratic perception

The strict searchlight analysis highlighted the angular gyrus as a region reflecting perceived similarity in both olfaction and vision. Such a representation is unlikely, however, to be confined to so restricted a locus. To ask whether the angular gyrus is indeed a hub for such representation, we measured its functional connectivity with 358 cortical parcels using beta-series correlations, averaged across participants and modalities. All 47 participants met the inclusion criteria. The cluster had a distributed connectivity profile encompassing default-mode and higher-order associative regions, including subgenual anterior cingulate and orbitofrontal cortex, frontopolar and ventrolateral prefrontal cortex, bilateral angular gyrus, precuneus, dorsolateral prefrontal cortex, and bilateral middle temporal cortex (Fig. 5a), consistent with its role as an associative-network hub.

**Figure 5.**
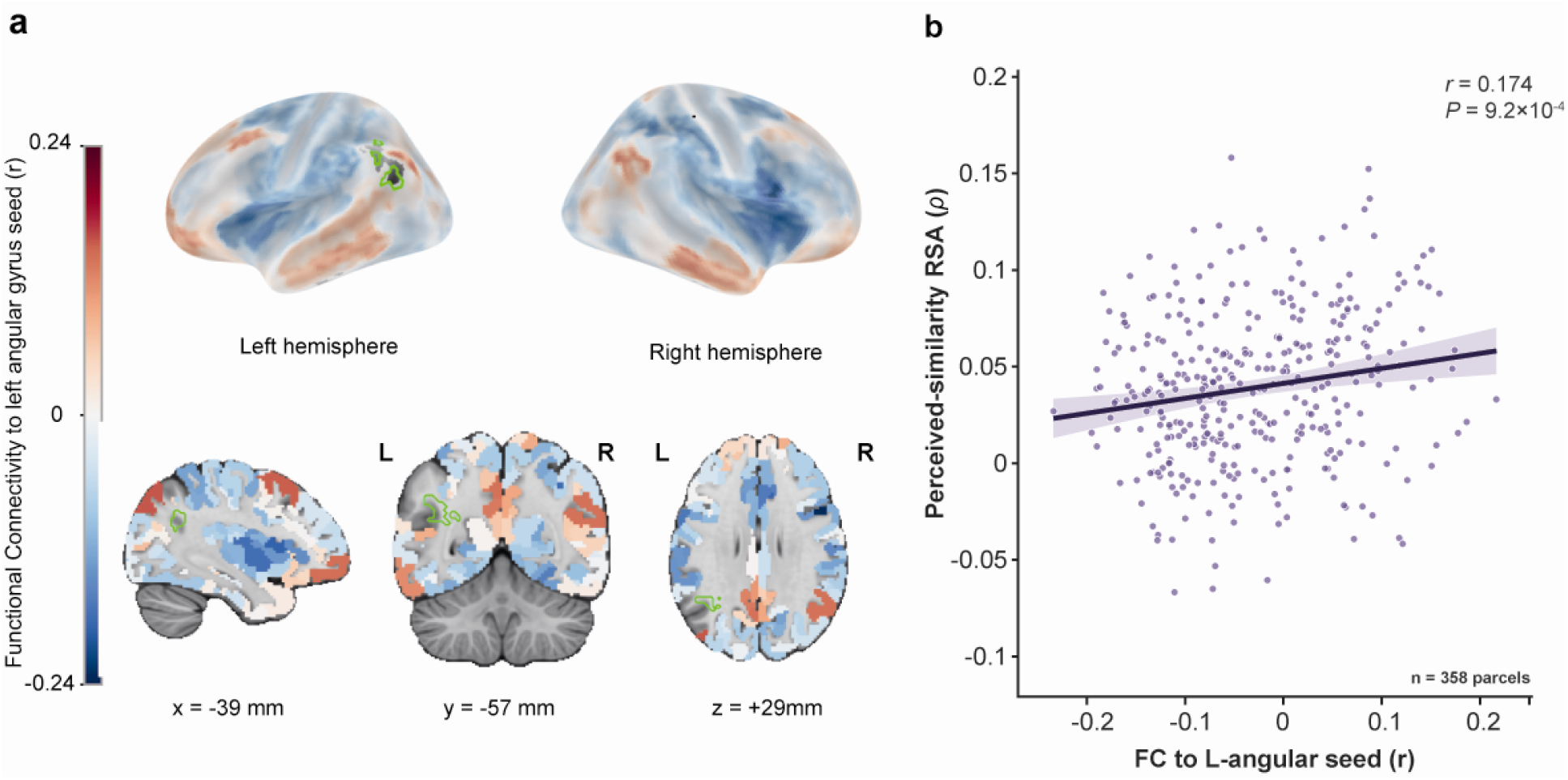
The left angular gyrus is the hub of a network reflecting idiosyncratic perception. **a,** Parcelwise functional connectivity to the cluster (beta-series FC, residualized against stimulus identity and global signal). **b,** Connectivity versus encoding across parcels. Each point is one cortical parcel (n = 358); x, its connectivity to the cluster (as in a); y, its perceived-similarity encoding (correlation between the parcel’s neural RDM and each participant’s own ratings, averaged across participants and modalities). Line, least-squares fit; shading, 95% CI.

We then asked whether regions more strongly coupled to the cluster also represented perceived similarity more strongly. For each parcel out of the Glasser HCP-MMP1 360-parcel atlas (n = 358, seed excluded), we related its connectivity with the cluster to its perceived-similarity RSA, and averaged across participants and modalities. Stronger connectivity was associated with stronger perceived-similarity encoding (combined Pearson *r* = 0.174 [95% CI: 0.072, 0.273], P = 9.2×10□□; Fig. 5b). This held in each modality separately (odor: *r* = 0.141 [95% CI: 0.038, 0.241], *P* = 0.0075; visual: *r* = 0.128 [95% CI: 0.024, 0.228], *P* = 0.016). This association was similar after controlling for parcel-to-seed distance (partial *r* = 0.165 [95% CI: 0.062, 0.264]), *P* = 0.002) and after excluding parcels within 20mm of the seed (n=352, *r* = 0.179 [95% CI: 0.076, 0.279], *P* = 7.2×10□□). The relationship also exceeded a spatial null distribution generated from 1,000 random rotations of the parcel centroids (null mean *r* = 0.030, SD = 0.076; *P* = 0.026). Thus, the angular gyrus is part of a network reflecting perceptual similarity, and given that it was the one locus that survived the strict thresholds of the previous whole-brain search, it can be viewed as its hub.

### The angular gyrus is centrally located in the path to idiosyncratic representation

Identifying the angular gyrus as a hub does not alone uncover its functional position in this network. To address this, we fitted deterministic, task-driven DCMs separately for olfaction and vision (N = 47), with the 221-voxel left AG cluster as a node (Fig. 4; Fig. 6). Driving inputs entered piriform cortex or V1/V2, respectively; endogenous connectivity was fully reciprocal, and group inference used parametric empirical Bayes with Bayesian model reduction (credible effects: Pp > 0.95).

**Fig. 6.**
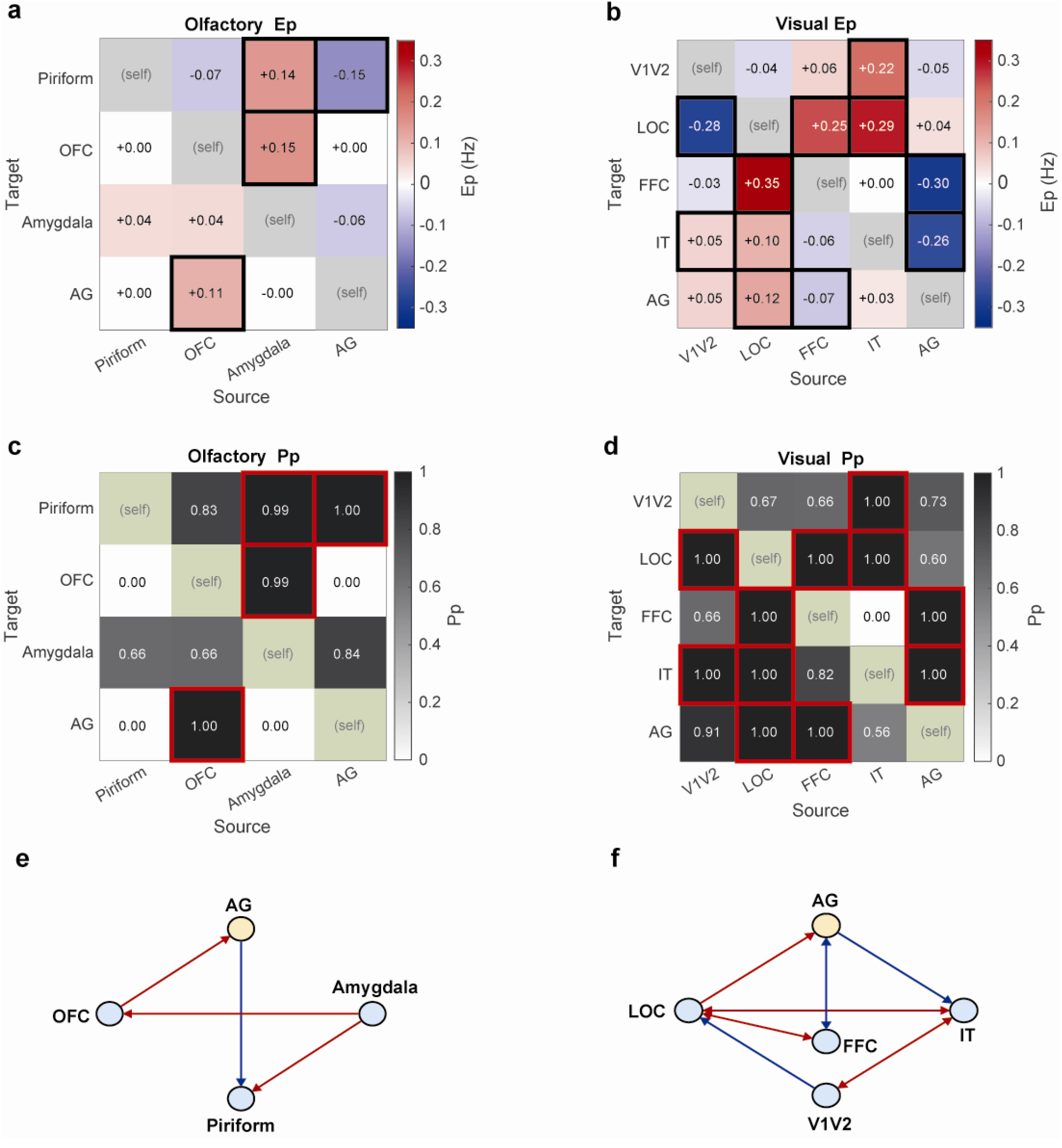
The angular gyrus is centrally located in the path to idiosyncratic representation. Deterministic single-state bilinear dynamic causal modelling (DCM) with parametric empirical Bayes and Bayesian model reduction (N = 47). Olfactory nodes: piriform, OFC, amygdala, AG; visual nodes: V1/V2, LOC, fusiform (FFC), IT, AG; stimulus drove the primary sensory node only. **a, b,** the effective-connectivity matrix (posterior expectation Ep, Hz) for olfaction (a) and vision (b). **c, d,** posterior probability (Pp) matrix for olfaction (c) and vision (d). In a-d, bold/red outline = credible (Pp > 0.95); grey diagonal = self-connections (not interpreted). **e, f,** layered graphs of credible directed connections for olfaction (e) and vision (f), arranged with the primary sensory node at the bottom and AG at the top, so that feedforward reads upward and feedback downward; double-headed arrows = reciprocal connections; color = sign of E⍰ (red +, blue −). The sign of an endogenous coupling is an effective rate constant defined relative to each region’s self-connection and does not denote synaptic excitation or inhibition; direction and posterior probability are the interpretable quantities.

In olfaction, the only credible feedforward input to AG came from orbitofrontal cortex (OFC → AG, E□ = 0.11 Hz, Pp = 1.00), not from piriform cortex or amygdala; AG in turn fed back to piriform (AG → piriform, E□ = -0.15 Hz, Pp = 1.00). The amygdala acted within the network, driving OFC and piriform (amygdala → OFC, E = 0.15 Hz; amygdala → piriform, E□ = 0.14 Hz; both Pp = 1.00), rather than coupling directly with AG. In vision, credible feedforward input to AG came from object-selective LOC (LOC → AG, E□ = 0.12 Hz) and fusiform cortex (FFC → AG, E□ = -0.07 Hz; both Pp = 1.00), while early visual cortex showed no credible coupling with AG in either direction (V1/V2 → AG, E□ = 0.05 Hz, Pp = 0.91; AG → V1/V2, E□ = -0.05 Hz, Pp = 0.73). V1/V2 influence was instead expressed within the visual hierarchy, as a credible projection to LOC (V1/V2 → LOC, E□ = -0.28 Hz, Pp = 1.00), consistent with a V1/V2 → LOC → AG relay. AG fed back on the ventral stream (AG → fusiform, E□ = -0.30 Hz; AG → IT, E□ = -0.26 Hz; both Pp = 1.00), fusiform thus being the only region reciprocally coupled with AG.

The models explained modest regional variance (group means 6.0% olfaction, 2.9% vision), so we emphasize the direction and credibility (Pp) of connections rather than their precise magnitude. Nor do we read coupling signs as synaptic excitation or inhibition: in bilinear DCM the sign denotes an effective rate constant relative to each region’s self-connection, and negative forward connections are common in this model class^33^. A common motif nonetheless emerged across modalities: a single region, OFC in olfaction and LOC in vision, provided the credible feedforward input to AG, while AG fed back on upstream sensory and limbic nodes, more extensively in vision (AG → fusiform, AG → IT) than in olfaction (AG → piriform). In other words, these findings depict the angular gyrus as a possible component in the emergence of the idiosyncratic representation, rather than merely a relay of a representation created elsewhere.

## Discussion

The world can be ordered by physical structure alone. Our internal ordering of that same world is shaped in addition by experience and makeup. Here we identify a cortical hierarchy that spans these two orderings. Modality-specific sensory cortices reflected the ordering given by the stimuli more strongly than the ordering participants reported. By contrast, the left angular gyrus reflected each participant’s own perceived ordering, in both olfaction and vision. Notably, an imaging study of the blue dress phenomenon noted in the introduction found that individual differences were reflected in the left inferior/superior parietal lobule^3^, which co-localizes with the angular-gyrus cluster identified here (Extended Data Fig. 5).

Previous studies asked these questions one at a time. In olfaction, some studies emphasized odor structure as the defining feature of the olfactory-cortex response^34,35^, and others emphasized odor perception^23,28,36^. In vision, some emphasized the physical properties of the stimulus^37,38^, and others object perception^39,40^. Here we contrasted both, in two senses, within the same individuals, and this yielded two observations. First, sensory cortex followed the physical models more closely than it followed perception. The brain therefore holds information about the stimulus that conscious report does not convey. Seconfviolid, a single region outside sensory cortex, the left angular gyrus, followed perception rather than physical structure, and did so in both senses. Connectivity analysis depicted this region as a possible hub for such representation, and dynamic causal modeling placed it downstream of modality-specific association cortex rather than of primary sensory areas^41,42^. Individually unique representational geometry has been described before within vision^43^, and we reproduce that result in inferior temporal cortex. What is new here is that one region carries it for two senses at once^44^. This matters because our odors and images shared no item identity, no semantic category and no physical feature space. The only thing common to them was relational geometry, the arrangement of stimuli relative to one another^9^. It is that arrangement, and not the stimuli themselves, that the angular gyrus holds in common across modalities.

Given the established role of the angular gyrus in conceptual and semantic processing^45,46^, one may raise the possibility that our results reflect a semantic mapping alone. Our data argue against this. Neither lexical-semantic similarity nor odor-identification ability explained the effect, and shared labels or taxonomic relations do not readily explain why neural structure followed each participant’s own odor judgements rather than the group consensus. The comparison of individual against consensus ratings is informative here. Visual similarity judgments were relatively consistent across participants, and angular-gyrus geometry was predicted equally well by an individual’s own ratings and by the group’s. Odor judgments were far more variable, and only a participant’s own ratings predicted their neural geometry; the group consensus did not. The angular gyrus therefore tracked perception as each person actually organized it, rather than perception as most people organize it. Rather than semantics, we suggest instead a representation upheld by the default-mode network, which combines incoming information with prior knowledge to construct context-dependent models of experience^47^. This fits the placement of the angular gyrus at the apex of a cortical gradient that extends from primary sensory areas to the default-mode regions^47,48^, and it fits lesion and stimulation evidence linking the angular gyrus to the integration of distributed information into coherent subjective and multimodal representations^46,49–54^. Our findings suggest that this integrative role extends beyond memory to the organization of individual perceptual experience.

Our study has several limitations. First, whereas the primary finding, a transition from stimulus-centered representations in modality-specific sensory cortex to a modality-general observer-centered representation in the angular gyrus, is, we think, very strong, the ensuing connectivity and directional analyses rest on inferences from imaging data that will require validation by other means. A future transcranial magnetic stimulation study of the angular gyrus, for example, could allow more definitive statements about its place in this hierarchy. Second, the stimulus sets were relatively small. This was dictated by the need for trial repetitions in the fMRI experiment combined with the long inter-stimulus intervals that olfaction demands, but it remains a limitation nonetheless. Third, we probed only two sensory systems, and extending these findings into added sensory domains may provide a more complete picture of the underlying brain organization. Finally, a semantic contribution to these results remains possible and deserves added attention. Despite these considerations, our findings reveal a cortical hierarchy in the organization of sensory perception. Modality-specific sensory systems preserved the ordering given by the stimuli, whereas the left angular gyrus preserved the ordering given by the observer, in both olfaction and vision. The brain thus maintains two accounts of the same external world, and our results place the angular gyrus at the transition between them. This suggests a potentially fundamental hierarchy in brain organization.

## Methods

### Participants

Fifty-two individuals participated in the fMRI experiment. Five were scanned with a preliminary visual stimulus set and were excluded from all analyses after two images were replaced following behavioral validation (see Stimulus-set construction and perceptual alignment). A separate panel of 30 individuals contributed to stimulus selection and was not scanned. The final sample comprised 47 participants (age 29.3 ± 7.08 years; 25 male, 22 female), all of whom completed an in-lab behavioral experiment (Experiment 1) followed by an fMRI experiment (Experiment 2; median interval, 8 days; IQR, 4–14 days). Sex was self-reported; the study was not designed or powered to test for sex differences, and sex was not included in any analysis. All participants reported no history of significant medical, psychiatric or olfactory dysfunction and provided written informed consent under a protocol approved by the ethics committee of Wolfson Medical Center (#0226-21-WMC). Participants were compensated at 100 NIS per hour for MRI scanning and 50 NIS for the one-hour behavioral session.

### Stimuli and physical models

Five monomolecular odorants were used: phenylethyl alcohol (PEA; PubChem CID 6054; CAS 60-12-8; 100%; rose-like), 3-cis-hexenol (CID 5281167; CAS 928-96-1; 1%; grass-like), isoamyl acetate (CID 31276; CAS 123-92-2; 1%; banana-like), methyl hexanoate (CID 7824; CAS 106-70-7; 10%; pineapple-like) and cuminaldehyde (CID 326; CAS 122-03-2; 10%; cumin-like). All odorants were obtained from Sigma-Aldrich, diluted in isopropyl myristate to the concentrations above, with concentrations selected to approximately match perceived intensity in pilot testing. Five images of real-world objects were used: basketball, strainer, filter, waterwheel and bookshelf, drawn from the THINGS database (Fig. 1).

For each modality we used an independently derived stimulus-computable measure of similarity, whose pairwise distances defined a model representational dissimilarity matrix (RDM). For odors we used the angle-distance model^10,11^ and for vision we used the SPoSE embedding^12–14^. Notably, both were developed against perceptual ratings obtained from human observers, yet we treat their outputs as physical measures. We can do this because what a model was built from and what it takes as input are separate matters. For example, the mapping from wavelength to perceived color was itself established psychophysically, through color-matching experiments in human observers, yet wavelength unarguably remains a physical quantity: it is obtained from the stimulus by measurement, with no observer in the loop. The same holds here. Once developed, the angle-distance metric returns a value computed from odorants alone^10,11^, and two recent studies have demonstrated that SPoSE returns a value computed from images alone^14,15^. For any given stimulus pair, the resulting value is therefore obtained from the stimulus rather than from a judgment, is fixed and identical across participants, and is independent of all behavioral and neural data collected here. In other words, we use the term physical because the value is stimulus-computable, in contrast to the idiosyncratic perceptual similarity reported by each individual.

### Stimulus-set construction and perceptual alignment

The stimulus set was developed in two stages. An initial set of five odorants and five objects was assembled by maximizing the correlation between the olfactory and visual physical-model RDMs, and five participants were scanned with this preliminary set; their behavioral ratings indicated weaker-than-expected correspondence between perceived odor and visual similarity. We therefore collected additional ratings from an independent behavioral panel (N = 30), who rated all pairwise similarities among an expanded candidate set of ten odorants and ten images on a 0–100 visual analog scale (two repetitions per pair). On the basis of these ratings alone, and before any analysis of neural data, two of the five images were replaced; the five preliminary participants were excluded from all subsequent analyses. From the panel-derived behavioral RDMs we defined the perceptual alignment as the one-to-one odor-object assignment (of 5! = 120 possible) that maximized correspondence between olfactory and visual perceived similarity (r = 0.62). This alignment matched relational similarity structure across modalities rather than semantic correspondence between individual odor-object pairs, and was fixed before recruitment of the final imaging sample (N = 47).

### Odor delivery

In the behavioral session, odorants were presented in sniff jars. In the scanner, odorants were delivered with a computer-controlled olfactometer and odor canopy, without nasal mask or cannula^55^. A Teflon nozzle ∼10 cm anterior to the participant’s nose delivered clean air at 1.5 l min□¹ with odor pulses embedded at trial onset, and continuous airflow through the head coil was maintained by a 2-inch vacuum hose. Nasal airflow was monitored with a spirometer (ML141, ADInstruments) and instrumentation amplifier (PowerLab 16SP, ADInstruments), enabling odor delivery to be triggered by inhalation onset.

### Experiment 1: In-lab behavioral experiment

Participants rated the perceived similarity of odor and image pairs in an in-lab behavioral experiment. For each modality, participants rated the similarity of all ten pairwise combinations of the five stimuli on a continuous 0–100 visual analog scale (0, no similarity; 100, maximal similarity), each pair three times, yielding 2,820 ratings. Ratings differing by >50 points from both other repetitions of the same pair were excluded (odor, 88 ratings, 6.25%; visual, 49 ratings, 3.48%), and one participant did not complete a subset of pairs, retaining 2,679. Odor pairs were presented in sniff jars and visual pairs on screen. The task comprised three runs, each containing all ten odor and all ten visual pairs with modalities alternating; each stimulus was shown for 1 s, stimuli within a pair separated by 3 s and successive pairs by 20 s. Before the odor task, participants rated each odor for pleasantness, intensity, familiarity and edibility and attempted to identify it (Extended Data Fig. 6); for images they named each object. For each participant and modality, ratings were converted to dissimilarities 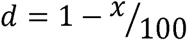 and arranged into a symmetric 5×5 RDM with zero diagonal.

### Behavioral analyses

To test whether ratings reflected the model-predicted structure, upper-triangular vectors of the group-mean behavioral RDMs were correlated with the corresponding physical-model RDMs across the ten unique pairs (Pearson), separately for each modality; significance was assessed with a one-sided t-test. Correspondence between the olfactory and visual group-mean behavioral RDMs was assessed identically. To visualize perceptual spaces, subject-level RDMs were embedded by classical two-dimensional multidimensional scaling and the olfactory configuration was aligned to the visual configuration by Procrustes analysis (rotation, reflection and isotropic scaling); significance was assessed by an exact permutation test over all 120 odor-image assignments, with P the proportion of assignments yielding a Procrustes distance ≤ the observed value. To quantify inter-individual consistency, we computed, for each participant and modality, the Spearman correlation between that participant’s RDM vector and the mean RDM vector of all other participants (leave-one-out consensus agreement). Agreement was tested against zero (one-sample t) and compared between modalities (paired t).

### Experiment 2: fMRI experiment

The same participants subsequently completed the fMRI experiment. MRI data were acquired using a 3T Siemens MAGNETOM Prisma scanner (Siemens Healthcare, Erlangen, Germany), equipped with a 32-channel receive head coil. Functional images were acquired with the multiband multi-echo gradient echo-EPI sequence^56,57^ using the following parameters: 63 axial slices oriented parallel to the anterior–posterior commissural (AC–PC) line, covering the whole brain. Four echoes were acquired with TEs = 10.60, 22.92, 35.24 and 47.56 ms; TR = 1,500 ms, flip angle = 69°; slice thickness of 2.4mm, no interslice gap, field of view = 216×216 mm²; and in-plane resolution of 2.4×2.4 mm^2^. A multiband acceleration factor of 3 was applied, parallel imaging (GRAPPA) factor of 3 (phase direction), and phase partial Fourier of 6/8. The multiband multi-echo EPI sequence was provided by the Center for Magnetic Resonance Research, University of Minnesota. T1-weighted anatomical images were acquired using a 3D T1-weighted Magnetization Prepared Rapid Gradient-echo (MP-RAGE) sequence: TR = 2,300 ms; TE = 2.29 ms; TI = 900 ms; flip angle = 8°; field of view = 240×240 mm²; matrix size = 256×256; 176 sagittal slices; voxel size = 0.94 mm isotropic; acceleration factor = 2.

During scanning, participants performed a 1-back task separately for each modality. In two olfactory runs each odorant was delivered six times per run (60 odor trials); in the visual run each image was presented 12 times (60 visual trials). Odor trials on which inhalation was not synchronized with odor delivery were excluded from all analyses (see Preprocessing and first-level modeling), so that only trials with delivery during inhalation contributed to pattern estimation. Participants pressed a button whenever the current stimulus matched the preceding one. No similarity ratings were collected in the scanner. Odor onset was sniff-triggered: the word “odor” appeared and the odorant was delivered for 3 s from inhalation onset (inter-trial interval jittered 17-20 s). Visual onset was likewise sniff-triggered: images were presented for 2 s (inter-trial interval jittered 13-16 s).

### fMRI data preprocessing and first-level modeling

Functional MRI data were analyzed using AFNI (version 25.1.15, “Maximinus”) together with custom MATLAB analysis scripts. Preprocessing was performed using afni_proc.py and included removal of the initial volumes, despiking, slice-timing correction using wsinc interpolation, motion correction, alignment to the anatomical image, nonlinear normalization to the MNI152 2009 template using SSwarper2, spatial smoothing (2 mm FWHM) within a brain mask, and voxel-wise scaling to percent signal change. Multi-echo data were optimally combined using TE-dependent denoising implemented in tedana (version 26.0.0)^58^. Volumes with framewise displacement > 0.8 mm or an outlier fraction > 10% were censored.

Trial-wise BOLD amplitudes were estimated with an individual-modulation GLM (3dDeconvolve, 3dREMLfit): each stimulus occurrence was modelled by a separate impulse-modulated regressor convolved with a canonical gamma HRF, giving one beta per trial, with REML estimation of temporal autocorrelation and motion parameters and censored time points as nuisance regressors. Because odor delivery was sniff-triggered, inhalation–delivery synchrony was verified on every odor trial from the simultaneously recorded nasal airflow. Odor trials on which inhalation was desynchronized from delivery (either no inhalation was detected during the delivery window or the odorant was delivered during exhalation) were excluded from all pattern and connectivity analyses; this removed 23 of 2,820 odor trials (0.8%) across 14 of 47 participants, with the corresponding trial-wise betas set to missing and omitted from all subsequent averaging. Visual trials were unaffected.

### Regions of interest and neural RDMs

ROIs were defined a priori from atlas-based masks registered to MNI space. HCP-MMP1 masks were used for V1/V2, LOC, FFC, IT; FreeSurfer Desikan-Killiany masks were used for OFC and amygdala; and piriform cortex frontal and temporal subdivisions were defined using dedicated anatomical masks from Zhou et al^59^. ROIs were restricted to voxels present in at least 80% of participants’ functional fields of view. The ROI sets comprised olfactory ROIs (OFC, amygdala, anterior and posterior piriform cortex)^23,59^, visual ROIs (V1/V2, LOC, FFC, IT)^26^.

For each participant, ROI and modality, multivoxel activity patterns were extracted from trial-wise beta estimates. Patterns were z-scored across voxels within each trial and then averaged across the retained repetitions of each stimulus. Neural dissimilarities were computed as Pearson correlation distance (1 - *r*) between stimulus-evoked multivoxel patterns, yielding one 5×5 neural RDM per participant, ROI and modality.

### ROI RSA analyses

For each participant and ROI, neural RDMs were correlated (Spearman) with predictor RDMs: the modality-matched physical model, each participant’s own behavioral RDM, and the group-consensus (mean) behavioral RDM; behavioral vectors were z-scored within participant and modality beforehand. Crossed predictor-to-neural analyses tested whether structure generalized across systems, correlating visual predictors with olfactory neural RDMs and odor predictors with visual neural RDMs. All RSA values were Fisher-z-transformed before group inference and back-transformed to mean Spearman *ρ* for reporting.

To test whether the stronger encoding of structural than perceptual similarity reflected differences in model reliability, we estimated within-participant perceptual-RDM reliability as the mean pairwise Spearman correlation across the three rating repetitions, adjusted to the full three-repetition estimate using the Spearman–Brown formula^60^. Perceptual RSA values were disattenuated by dividing by √R; the fixed structural model was assigned R = 1. We also computed the Nili noise ceiling for each ROI and normalized RSA values accordingly^8^. Corrections used the group-median reliability for each modality; participant-specific correction, excluding degenerate estimates (R ≤ 0.1), yielded the same conclusions.

### Searchlight RSA

Whole-brain searchlights used 8 mm-radius spheres (minimum 30 voxels). At each center, trial-wise beta patterns were processed as in the ROI analysis and converted to a 5×5 neural RDM by correlation distance, then correlated (Spearman) with that participant’s own behavioral RDM, yielding one map per participant and modality. Group analyses were performed at two levels. First, each modality was analyzed separately: subject-level maps were Fisher-z-transformed and submitted to group one-sample tests against zero, corrected both by permutation-based cluster-level family-wise error control (voxel-wise P < 0.001; 2,000 sign-flip permutations; NN = 1 face-adjacent connectivity; cluster P < 0.05) and, as a complementary cluster-extent–independent control, by Storey’s adaptive positive false-discovery-rate procedure (q < 0.05)^61^. Second, a combined perceived-similarity map was formed by averaging the two single-modality maps and tested identically against zero, with cluster-level FWE correction only. A 6 mm-radius searchlight was run identically as a sensitivity analysis. Mean RSA values extracted from surviving clusters were used only for effect-size visualization, split-half robustness and the follow-up characterizations below; no further inference was performed on the extracted values themselves.

### Angular-gyrus cluster analyses

For the left angular-gyrus cluster, participant-level mean RSA was tested against zero separately for olfaction, vision and their combination. Because modality-specific effects were extracted from a cluster defined using the combined map, they were validated using leave-one-subject-out cross-validation. For each held-out participant, the cluster was redefined from the remaining 46 participants as the 100 voxels with the highest combined RSA within an anatomically constrained angular-gyrus search region, and the held-out participant’s mean RSA was extracted from those voxels. Results were similar for cluster sizes of K = 50, 100 and 221 voxels. Spatial specificity was assessed against 1,000 size-matched random-voxel clusters sampled from the group EPI-coverage mask, and robustness was evaluated across 1,000 split-half resamples. Robustness to individual stimulus pairs was assessed using a leave one pair out analysis. Each of the ten unique stimulus pairs was removed in turn from both the neural and behavioral RDM vectors, and Spearman RSA was recomputed across the remaining nine pairs. The resulting values were Fisher z transformed, averaged across participants and tested against zero using one-sided tests.

We next tested whether the cluster represented participant-specific perceived similarity rather than stimulus-derived or shared structure. Modality-matched physical-model RDMs were correlated with the cluster neural RDMs, and partial correlations quantified perceived similarity controlling for the physical model, and vice versa. Partial-correlation estimates were Fisher-z transformed, averaged across modalities and tested against zero. Participant-specific behavioral RDMs were then replaced with leave-one-out group-consensus RDMs, and individual and consensus effects were compared (paired t). To test the dissociation between sensory cortex and the angular gyrus, we computed per participant the physical-minus-perceptual RSA difference (Fisher-z) in the a priori sensory ROIs and in the angular gyrus, and evaluated the Region × Predictor interaction as a one-sample t-test on [(sensory physical − sensory perceptual) − (AG physical − AG perceptual)].

To assess semantic mediation, combined RSA was correlated with general and specific odor-identification performance; visual identification was at ceiling. We also constructed lexical-semantic RDMs as 1 – Wu-Palmer similarity (WordNet)^32^ between the stimulus labels (odors: rose, grass, banana, pineapple, cumin; the five object names for images) and tested their correspondence with cluster neural and perceived-similarity RDMs. Odor-identification and semantic analyses were two-sided; all other cluster tests were one-sided. Follow-up analyses were FDR-corrected within four prespecified families: perceptual RSA, physical-model RSA, perceived-versus-model partial correlations and odor-identification correlations. To test whether the cluster effect reduced to low-level odor attributes, we built per-participant dissimilarity matrices from absolute pairwise differences in rated pleasantness and intensity and computed partial Spearman correlations between the cluster neural RDM and the perceived-similarity RDM controlling for these attributes (N = 44 with complete ratings).

### Seed-based connectivity and connectivity-encoding analysis

The angular-gyrus cluster served as a seed for beta-series functional connectivity. Target parcels were the HCP-MMP1 360-parcel atlas restricted to voxels covered in ≥80% of participants and ≥20 voxels; parcels overlapping the seed (bilateral PGi) were removed, leaving 358 parcels. Per participant and modality, seed and parcel mean trial-wise beta series (60 trials) were jointly residualized against five stimulus-identity regressors and the global signal within the group EPI-coverage mask, and functional connectivity was computed as the Pearson correlation of the residual series; connectivity was Fisher-z-transformed and averaged across participants and then across modalities. Participants were retained per modality if ≥40 trials survived, with ≥2 per stimulus; all 47 participants met criterion. To test whether more strongly coupled regions encoded perceived similarity more strongly, each parcel’s connectivity to the seed was correlated across parcels (Pearson) with its combined perceived-similarity RSA. Robustness was assessed by (i) partial correlation controlling for parcel-to-seed Euclidean distance, (ii) exclusion of parcels within 20 mm of the seed, and (iii) a spatial-autocorrelation null generated by 1,000 rigid rotations of the parcel centroids with nearest-parcel reassignment, recomputing the across-parcel correlation each time (two-sided P as the proportion of |r| ≥ observed).

### Dynamic causal modelling

Directed connectivity was estimated with deterministic, single-state DCM^33^ (bilinear, with no modulatory [B] terms) in SPM12 (N = 47). Each modality was modelled as a single network containing the 221-voxel left angular gyrus (AG) searchlight cluster: an olfactory model (piriform cortex, OFC, amygdala, AG) and a visual model (V1/V2, LOC, fusiform [FFC], IT, AG). Node time series were the first eigenvariate of each ROI; for the anatomical ROIs (FreeSurfer OFC and amygdala; Glasser V1/V2, LO1–LO2, FFC, TE2a/b; bilateral piriform), voxels were restricted to those exceeding the whole-model F threshold (p < 0.05, uncorrected), with any ROI retaining < 15 such voxels reverting to its full mask; AG was taken whole, without further voxel selection. Eigenvariates were extracted from preprocessed BOLD after regressing out motion parameters, their derivatives, and per-run polynomial drifts (≤ 3rd order). A single lumped input (3 s [odor] or 2 s [visual] boxcars at all stimulus onsets) entered the primary sensory node only; endogenous (A) connections were fully reciprocal. Subject-level models were inverted, then combined with parametric empirical Bayes, followed by Bayesian model reduction and averaging over the A matrix (single group-mean regressor). Connections were credible at posterior probability (Pp) > 0.95.

### Statistical inference

ROI RSA scores were Fisher-z-transformed and tested against zero with one-sample t-tests; directional (one-sided) tests were used for planned positive RSA effects, and effect sizes are reported as Cohen’s *dz*. Semantic-encoding, odor-identification, connectivity-encoding, and spatial-null (spin) tests were two-sided; all other RSA, partial, LOO tests were one-sided (directional positive). Group means are reported as mean Spearman ρ after back-transformation. To compare physical-model and reported-perceptual effects across systems, a 2×2 repeated-measures ANOVA was run on Fisher-z RSA values averaged across the four ROIs of each modality, with within-participant factors Region-Type (olfactory, visual) and Similarity-Measure (physical model, own ratings), and partial η² as the effect-size measure; follow-up paired t-tests compared the two measures within each Region-Type. FDR correction (Benjamini–Hochberg) was applied within each modality across eight planned tests (four ROIs × [physical model or own ratings]); group-consensus behavioral analyses and crossed predictor-to-neural analyses were corrected separately within each modality. Task performance was quantified as d′ (log-linear corrected), tested against zero with two-tailed one-sample t-tests in each modality. Results are reported as qFDR, with qFDR < 0.05 considered significant.

## Acknowledgements

We would like to thank Prof. Rafi Malach, Dr. Shani Grossman and Prof. Galit Yovel for their important comments and suggestions.

## Data Availability

Behavioral data are available at the git repository: https://gitlab.com/michal.andelman/odor-visual-rsa. Neural data are available at OpenNeuro link: https://openneuro.org/datasets/ds008255.

## Code Availability

All code is available at the git repository: https://gitlab.com/michal.andelman/odor-visual-rsa.

## Author contributions

Conceived idea: M.A.G., N.S.; Designed experiments: M.A.G., T.W., O.P., N.S.; Recruited participants: M.A.G., T.W., Programmed the olfactometer experiment: D.H., L.G., M.A.G; Conducted experiments: M.A.G., T.W.; Analyzed data: M.A.G., N.S.; Wrote first draft: M.A.G.; Edited final draft: All authors.

