## Extended Data for "A cortical hierarchy of sensory representations from physical structure to idiosyncratic perception": Extended Data.docx

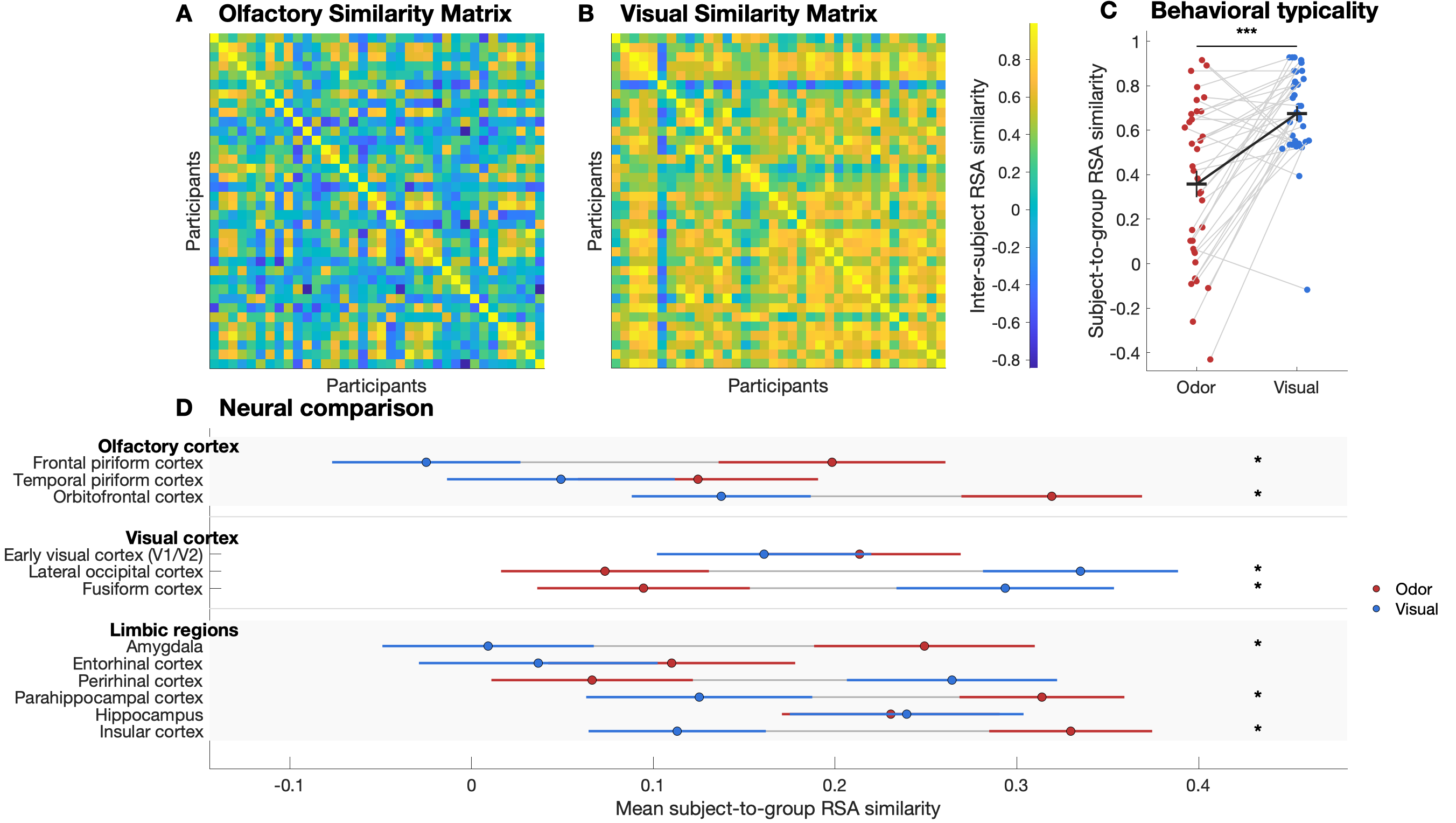

**Extended Data Fig 1: Subjects were more similar in visual ratings than olfactory ratings**

**a,b,** similarity matrices between participants for olfactory (a) and visual (b) ratings. **c**, subject to group RSA similarity. Each circle represents a participant. ****p* < 0.001.

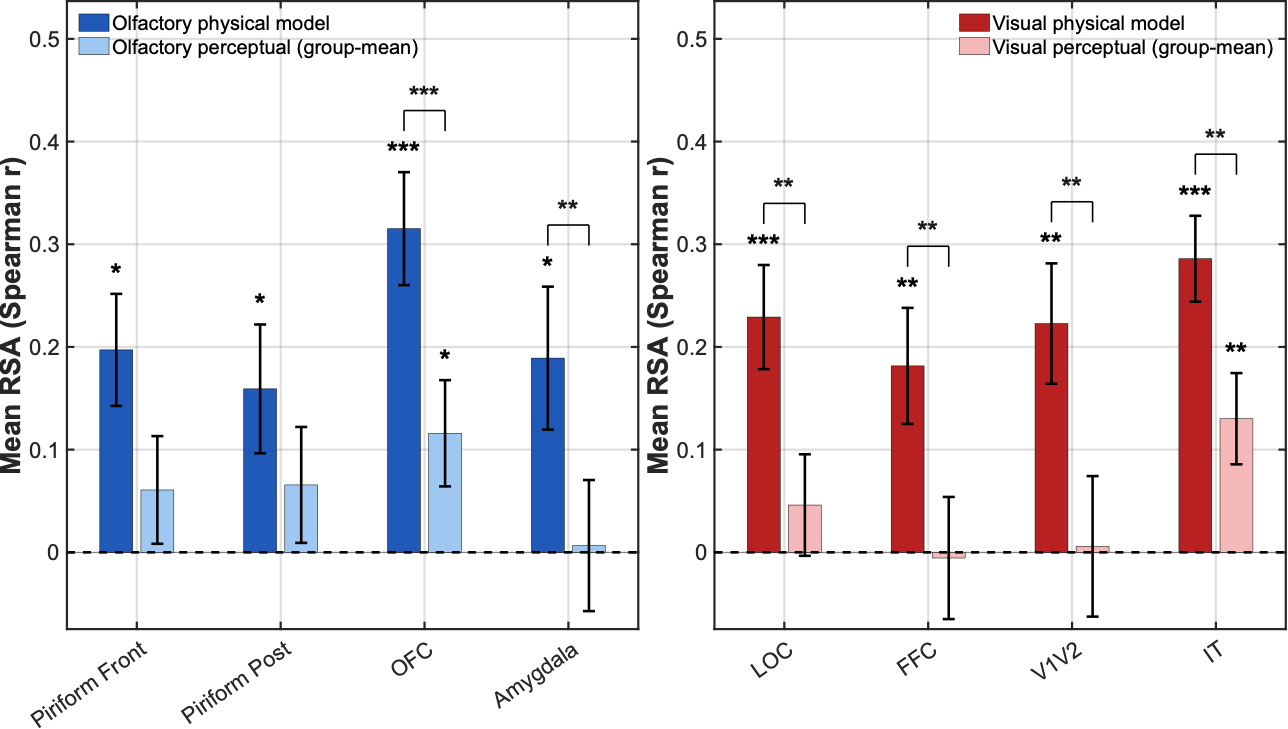
.

**Extended Data Fig. 2: in sensory cortices, within-modality physical RSA was stronger than group-mean perceptual RSA**

Left: bars present mean RSA between olfactory neural RDMs and either the olfactory physical RDM (dark blue) or the olfactory group-mean perceptual RDM (light blue) across olfactory ROIs. Right: bars present mean RSA between visual neural RDMs and either the visual physical RDM (dark red) or the visual group-mean perceptual RDM (light red) across visual ROIs. Bars indicate group mean and error bars indicate SEM. Stars denote FDR-corrected significance across ROIs within each planned analysis family: **q*FDR < 0.05, ***q*FDR < 0.01, ****q*FDR < 0.001.

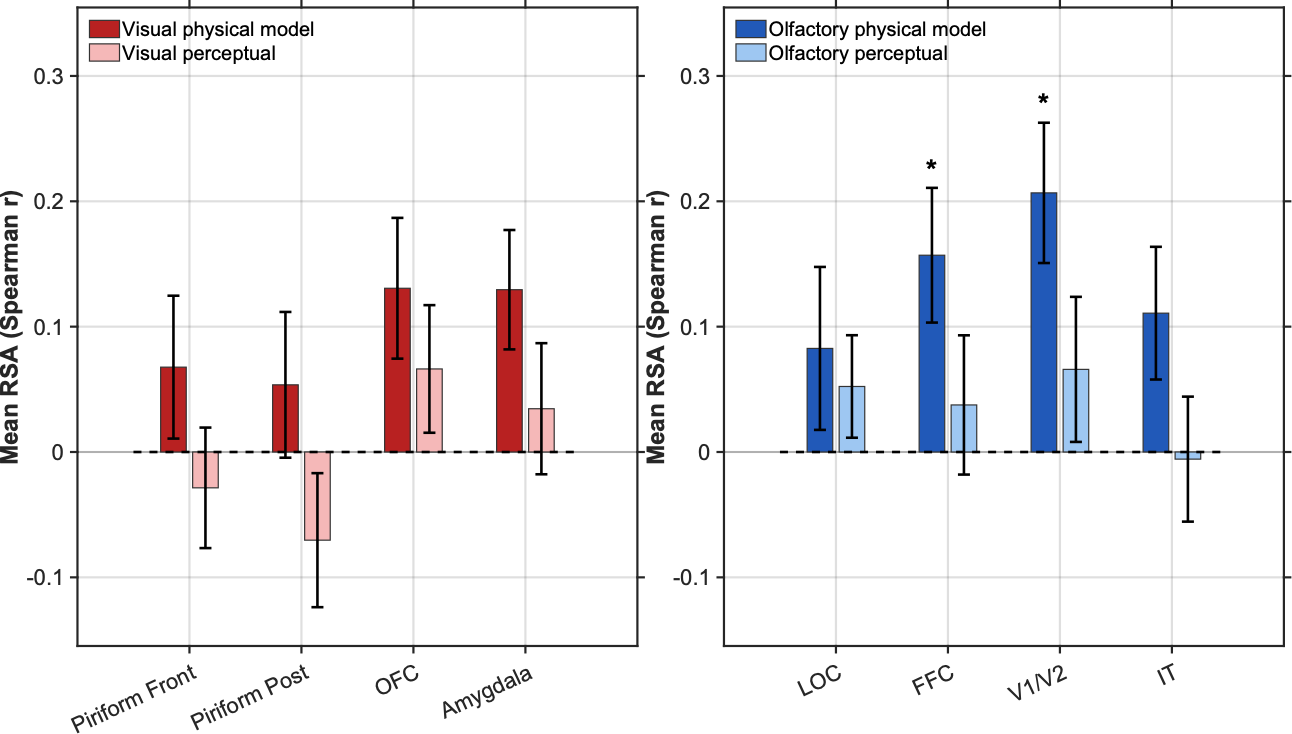

**Extended Data Fig. 3: Crossed physical-to-neural RSA revealed odor-physical effects in visual cortex but no visual-physical effects in olfactory cortex**

Left: bars present mean RSA between olfactory neural RDMs and either the visual physical RDM (dark red) or the mean-group visual perceptual RDM (light red) across olfactory ROIs. Right: bars present mean RSA between visual neural RDMs and either the olfactory physical RDM (dark blue) or the olfactory group-mean perceptual RDM (light blue) across visual ROIs. Bars indicate group mean and error bars indicate SEM. Stars denote FDR-corrected significance across ROIs within each planned analysis family: **q*FDR < 0.05, ***q*FDR < 0.01, ****q*FDR < 0.001.

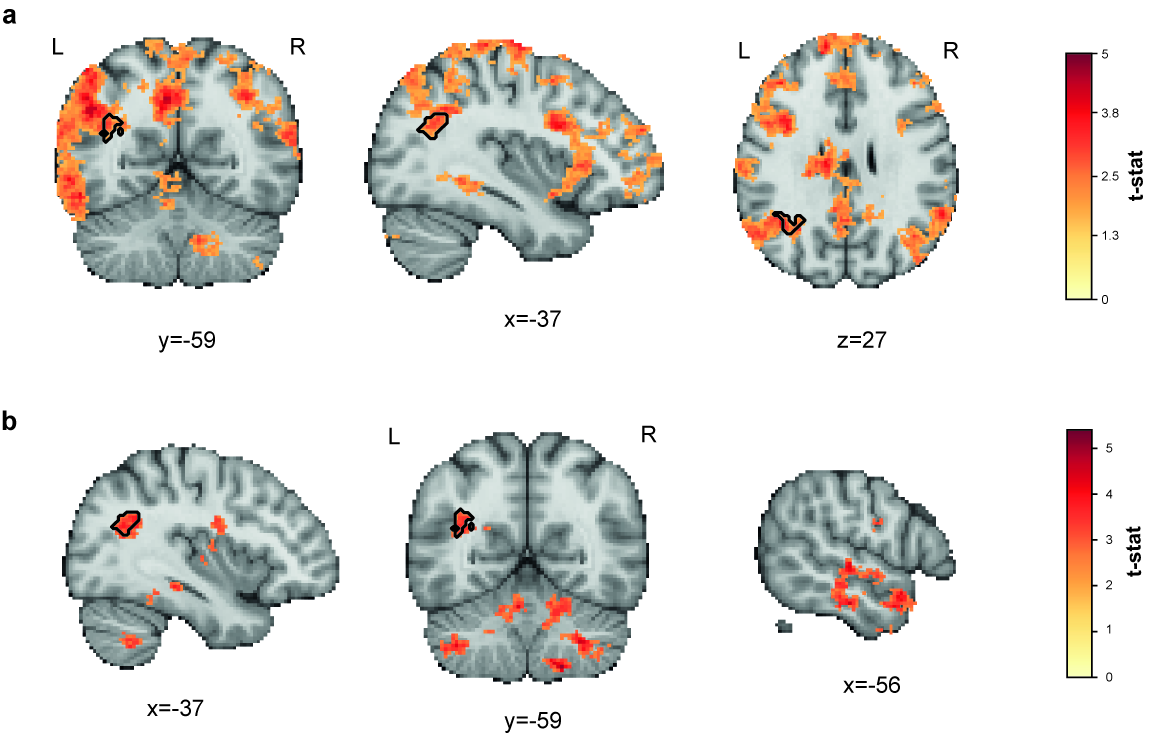

**Extended Data Fig. 4: Whole-Brain perceptual RSA overlapped in the angular gyrus across vision and olfaction**

**a**,**b**, Whole-brain searchlight maps for odor (a) and vision (b). At each 8-mm-radius sphere, the local neural RDM was correlated with each participant’s own perceptual-similarity RDM and tested against zero across participants (N = 47). Maps present voxels surviving adaptive Storey FDR correction at q < 0.05. Color indicates the group t-statistic. The black outline marks the largest odor-vision overlap cluster in the left angular gyrus (132 voxels). For display only, clusters (using 6-connectivity) smaller than 100 voxels were removed. Sections pass through the angular gyrus (a: x = −37, y = −59, z = 27; b: x = −37, y = −59) and, for vision, the occipitotemporal cluster (x = −56).

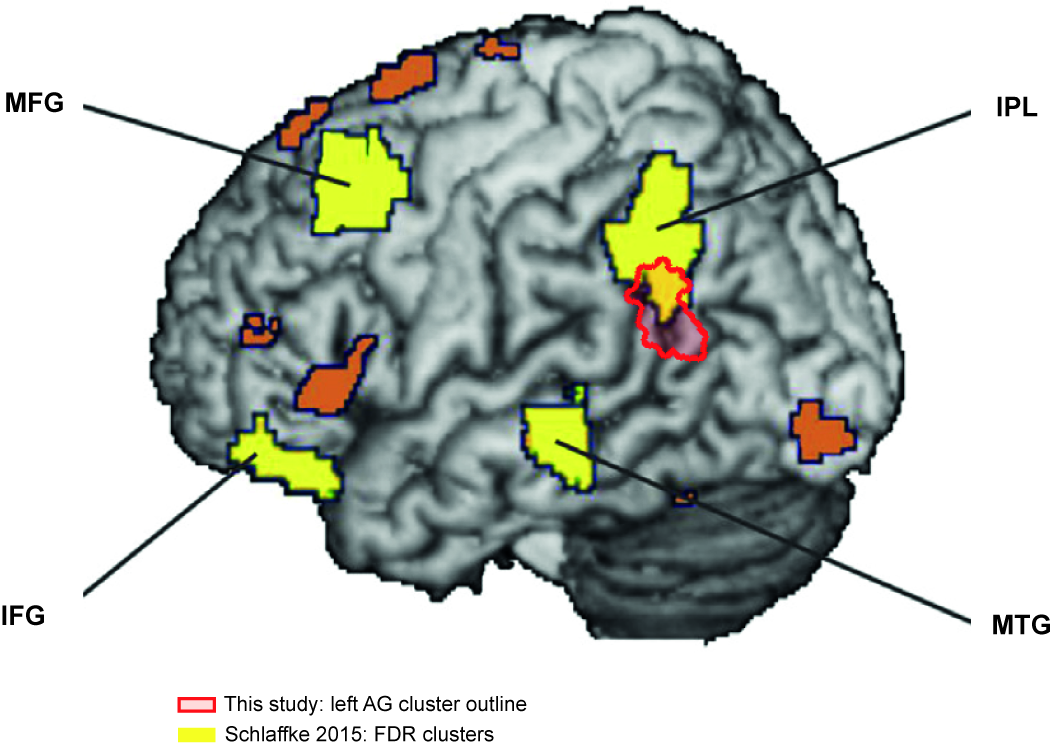

**Extended Data Fig. 5: The angular-gyrus cluster co-localized with a left parietal region associated with perception of “The Dress”**

Our left angular-gyrus cluster (red outline; 221 voxels) overlaid on Fig. 3 of Schlaffke et al. (2015, Cortex), which reported greater responses to the dress in white/gold than blue/black perceivers (yellow = FDR-corrected clusters; orange = additional, uncorrected). As only their published figure and peaks were available, our cluster was projected onto their render using their four clusters as fiducials. It overlaps the inferior aspect of their left SPL/IPL cluster (peak -46, -54, 43;) - the same inferior-parietal territory.

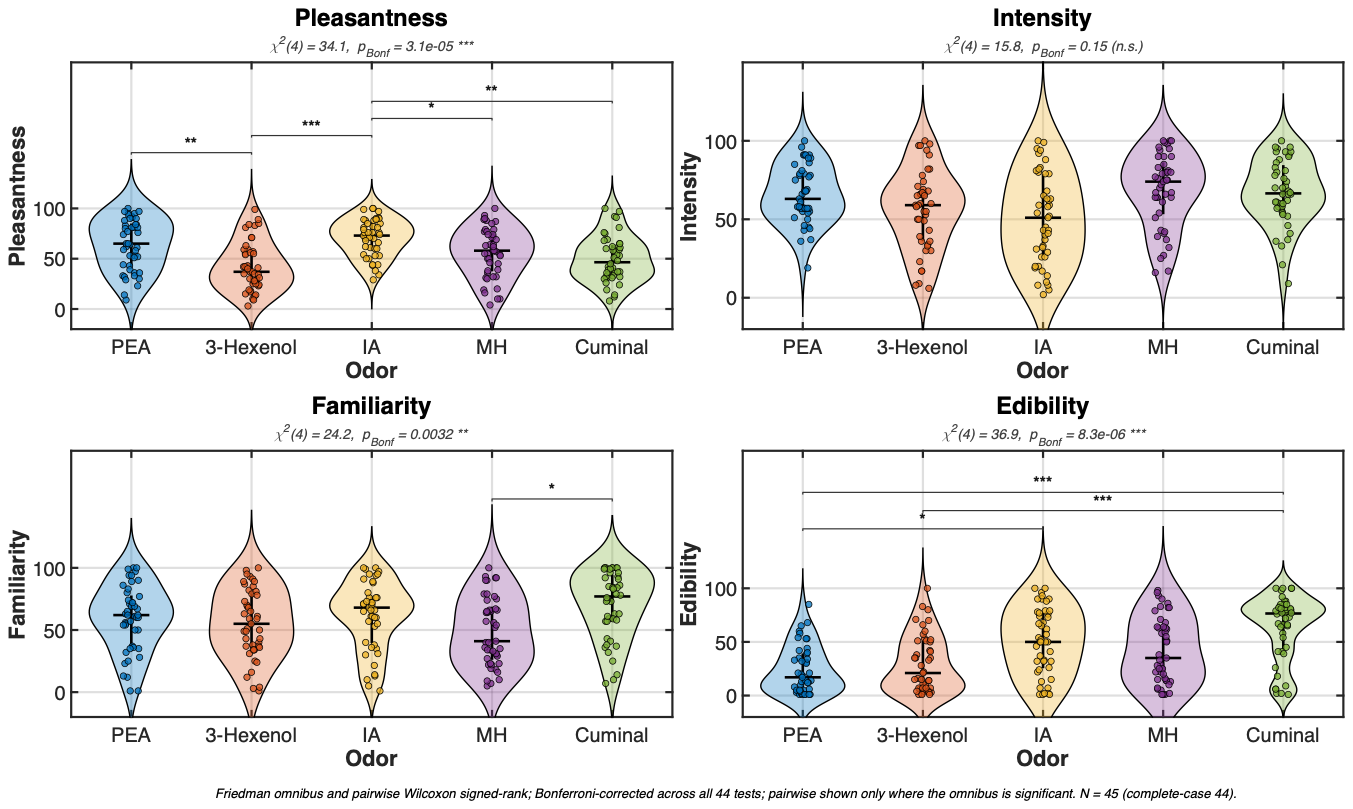

**Extended Data Fig. 6: Odors were iso-intense, but ranged in pleasantness and edibility**

Violin plots presenting odorant (blue = PEA, red = 3-Hexenol, yellow = Isoamyl Acetate, purple = Methyl Hexanoate, green = Cuminaldehyde) ratings along perceptual descriptors (pleasantness, intensity, familiarity, edibility). *P*-values are indicated. Dots represent individual participants; the horizontal line and vertical bar indicate the median and interquartile range; the violin represents the distribution. N = 45.

**Extended Data Table 1 | Disattenuation analysis of physical and perceptual RSA**

| ROI | Modality | Physical *ρ* | Perceptual *ρ* (observed) | Perceptual *ρ* (disattenuated) | Physical vs. disattenuated perceptual, *P* |
| --- | --- | --- | --- | --- | --- |
| Orbitofrontal | Olfactory | 0.314 | 0.094 | 0.134 | 0.026 |
| Anterior piriform | Olfactory | 0.200 | 0.156 | 0.231 | 0.66 |
| Posterior piriform | Olfactory | 0.150 | 0.148 | 0.198 | 0.54 |
| Amygdala | Olfactory | 0.186 | 0.018 | 0.017 | 0.05 |
| Inferior temporal | Visual | 0.287 | 0.135 | 0.148 | 0.037 |
| Lateral occipital | Visual | 0.223 | 0.042 | 0.044 | 0.009 |
| V1/V2 | Visual | 0.224 | −0.011 | −0.010 | 0.007 |
| Fusiform (FFC) | Visual | 0.185 | 0.042 | 0.045 | 0.049 |

For each ROI, values show group-mean RSA (Fisher-z-averaged Spearman *ρ*; N = 47) with the physical model and each participant’s perceptual RDM. Perceptual RSA was disattenuated by dividing by √R, using Spearman–Brown-corrected median reliabilities of R = 0.71 for odors and R = 0.88 for images; the fixed physical model was assigned R = 1. The final column reports two-sided paired t-tests on Fisher-z values comparing physical and disattenuated perceptual RSA.

Because the physical model was fixed across participants and was assigned reliability R = 1 whereas perceptual ratings are reliability-limited (Spearman-Brown: odor *R* =0.71; vision *R* = 0.88), we tested whether this difference explained the physical-model advantage. After deattenuating perceptual RSA for measurement reliability24, the advantage remained in visual cortex (physical *ρ* = 0.230 versus corrected perceptual *ρ* = 0.057; *t*(46) = 3.51, *P* = 0.001) and across all visual ROIs. In olfactory cortex, it also persisted in OFC and amygdala, whereas in piriform cortex, where perceptual similarity was significantly encoded, the two models became comparable after correction (anterior piriform: 0.2 vs. 0.23; posterior piriform: 0.15 vs. 0.198). Thus, differences in perceptual-RDM reliability did not explain the physical-model advantage in visual cortex or in OFC and amygdala. In anterior and posterior piriform cortex, however, reliability correction rendered physical and perceptual correspondence statistically comparable.
